# JNK regulates β3-containing GABAAR expression at the cell surface via the receptor clustering protein GIT1 (ArfGAP1)

**DOI:** 10.1101/2025.01.22.634236

**Authors:** Artemis Varidaki, Prasannakumar Deshpande, Elina Nagaeva, Annika Schäfer, Christel Sourander, Ye Hong, Soroush Abyari, Valentina Siino, Christoffer Lagerholm, Peter James, Esa R. Korpi, Eleanor Coffey

**Affiliations:** Turku Bioscience Centre, University of Turku and Åbo Akademi University, Turku 20520, Finland; Department of Pharmacology, Medicum, Faculty of Medicine, University of Helsinki, 00014 Helsinki, Finland; Department of Immuntechnology, Scheelevägen 2, Medicon Village, House 406, Lund University, Sweden

**Author notes:** Corresponding author: Eleanor Coffey, Turku Bioscience Centre, University of Turku and Åbo Akademi University, FIN-20520, Finland. Equal contribution (Artemis Varidaki, Prasannakumar Deshpande). The authors declare no conflicts of interest either financial or other kind. UK Dementia Research Institute at Imperial College London, Department of Brain Sciences, London W12 0BZ, United Kingdom.

## Abstract

GABA_A_-type receptors (GABAARs) mediate fast and tonic inhibition and are essential for maintaining excitatory/inhibitory balance in neural circuits. Disrupted GABAergic signaling leads to maladaptive changes associated with neuropsychiatric disorders, yet the mechanisms that regulate GABAAR surface availability remain incompletely understood. We previously identified that the stress-activated kinase JNK1 drives anxiety-like behaviours in mice suggesting a potential link between stress signalling and GABAergic regulation. Here, we show that genetic deletion or inhibition of JNK1 increases surface expression of β3-containing GABAARs at extrasynaptic sites and at excitatory synapses where they may participate in shunting inhibition. We identify the signalling scaffold GIT1 as a direct JNK1 substrate, phosphorylated at S371, and find that GIT1 accumulates in dendritic spines in *Jnk1-/-* neurons or following JNK inhibition, whereas the phosphomimetic GIT1-S371D is excluded. Moreover, GIT1 is required for JNK-dependent regulation of β3-GABAAR trafficking. Accordingly, neurons from *Jnk1-/-* mice exhibit increased spontaneous inhibitory postsynaptic potentials and enhanced tonic inhibition. These results reveal a JNK1-GIT1 signalling axis that suppresses GABAAR stability and inhibitory tone, providing a mechanism by which the stress-activated JNK pathway modulates synaptic and extrasynaptic inhibition.

## INTRODUCTION

Gamma-aminobutyric acid (GABA) is the main inhibitory neurotransmitter in brain. It serves an important role to balance excitatory neurotransmission, whereas deficit in GABAergic function is associated with a range of psychiatric disorders, in particular major depression and schizophrenia (Chang et al., 2003; Rubenstein and Merzenich, 2003; Hasler et al., 2007; Eichler and Meier, 2008; Ramocki and Zoghbi, 2008; Uhlhaas and Singer, 2010; Luscher et al., 2011a; Harrison, 2015). In particular, GABAARs play a key role in modifying anxiety and benzodiazepines, which increase GABAAR opening, are widely used for treatment of anxiety disorders (Kalueff and Nutt, 2007).

GABAergic deficit can result from reduced GAD67 expression leading to lower GABA synthesis (Lewis et al., 2012; de Jonge et al., 2017), or from loss of GABAergic interneurons themselves, leading to lower GABA and elevated hippocampal activity (Cohen et al., 2015; Heckers and Konradi, 2015). However, GABAergic deficit is also associated with receptor trafficking defects (Mueller et al., 2015; Wei et al., 2015; Mele et al., 2019). Notably, GABAAR response size directly correlates with the number of available receptors at the cell surface (Jacob et al., 2008; Luscher et al., 2011b), indicating that this is the rate limiting aspect of GABAergic function, and even small changes in receptor levels have been shown to affect behaviour in mice (Nusser et al., 2001; Vicini and Ortinski, 2004).

GABAARs are pentameric ligand gated Cl^-^ channels, consisting of diverse compositions of α, β and γ subunits. These receptors mediate phasic inhibitory synaptic transmission and tonic inhibitory currents from extrasynaptic and perisynaptic locations, leading to long lasting inhibition that controls network activity (Olsen and Sieghart; Schipper et al., 2016). However, GABAARs have also been observed at glutamatergic synapses. Electron microscopy studies have shown GABAAR β2 and β3 subunit expression in the post-synaptic density of asymmetric synapses, coexisting with glutamate receptors and consistent with co-release of both neurotransmitters (Nusser et al., 1998; Fattorini et al., 2009; Marlin and Carter, 2014). Also, in cortex and in hippocampus, there is evidence for mixed synapses where VGAT and VGLUT1 are co-expressed in mixed synapses (Safiulina et al., 2006; Zander et al., 2010; Fattorini et al., 2015). However, studies addressing the mechanisms that govern the stabilization of GABAARs have not taken into account the pool of receptors that localise at excitatory synapses. Several molecules are known that control trafficking and stabilization of GABAARs to and from the cell surface, and subsequent stabilization at inhibitory synapses (Lorenz-Guertin and Jacob, 2018). One of the recently identified regulators of GABAAR clustering is a multi-domain protein known as G-protein coupled receptor kinase interacting protein-1 (GIT1) (Smith et al., 2014).

GIT1 is ubiquitously expressed in brain where it is found both pre- and post-synaptically (Kim et al., 2003; Smith et al., 2014; Zhou et al., 2016). Also known as Arf GAP-1, GIT1 harbours an Arf-regulating GAP domain; a Spa2 homology domain (SHD) through which it interacts with PAK-interacting exchange factor-β (β -PIX) and a C-terminal paxillin binding domain (PDB) (Zhao et al., 2000; Schlenker and Rittinger, 2009; Zhou et al., 2016). GIT1 knockout mice exhibit learning deficits and reduced hippocampal spine density (Schmalzigaug et al., 2009; Menon et al., 2010; Hong and Mah, 2015; Martyn et al., 2018; Fass et al., 2022), while knockout of its homolog GIT2 produces an anxiogenic phenotype (Schmalzigaug et al., 2009). A single nucleotide variant in GIT1 is associated with attention deficit hyperactivity disorder in humans and with hyperactivity in mice (Won et al., 2011), whereas 37 single nucleotide variants in GIT1 were identified in a schizophrenia cohort, including function disrupting missense mutations (Kim et al., 2017a). Also, GIT1 mRNA is decreased in the hippocampus of depressed suicide victims (Fuchsova et al., 2015). Consistent with these associations of *GIT1* anomalies with neuropsychiatric dysfunction, *Git1^-/-^* mice show altered excitatory/inhibitory balance with reduced inhibitory neurotransmission (Won et al., 2011). Together these studies suggest that GIT1 serves an important role in excitatory/inhibitory homeostasis.

The highly conserved c-Jun N-terminal kinase (JNK) pathway acts as a sensor of extracellular stimuli, triggering adaptive cellular responses (Hotamisligil and Davis, 2016). In the nervous system, JNK regulates anxiety and depressive-like behaviours in mice (Mohammad et al., 2018; Stefanoska et al., 2018), while genetic anomalies on the JNK pathway are associated with schizophrenia (Winchester et al., 2012; Winchester et al., 2014; Marchisella et al., 2016). Yet, the molecular mechanism whereby JNK contributes to such disorders is not known. Here, we identify that physiologically active JNK phosphorylates GIT1 on S371 (S392 in human). We demonstrate that the JNK1 phosphorylation site on GIT1 controls its translocation to spines and is required for GABAAR stabilization at the cell surface in dendritic spines and in dendrites. Moreover, *Jnk1* deletion increases GABAAR colocalization with VGLUT1-positive synapses in spines, and with VGAT-positive synapses in shafts, indicating that JNK1 regulates GABAAR presence at excitatory synaptic sites, where it may play a modulatory role, and at extrasynaptic sites. Consistent with this, amplitudes of spontaneous inhibitory post-synaptic potentials (sIPSCs) and tonic GABA current are enhanced in *Jnk1-/-* neurons. These findings suggest that JNK1 can control cell surface GABAAR stability at synaptic and extrasynaptic sites via a mechanism involving receptor-clustering scaffold GIT1.

## MATERIALS AND METHODS

### Source of plasmids

pEGFP-C1 GIT1 (EGFP GIT1 WT) (Addgene plasmid: # 15226) and Flag-beta PIXα (Addgene plasmid: # 15234) were gifts from Rick Horwitz. pSuper-GIT1 small hairpin RNA (shRNA) was a generous gift from Huaye Zhang. For in vitro phosphorylation assay, pEBG-JNK1a1 and pEGFP-MEKK1Δ(1174-1493) has been previously described (Björkblom et al., 2008). pmRuby2-Lifeact7 was from Michael Davidson’s lab via Addgene, pEGFP-Dynamin 2-WT and pEGFP-Dynamin 2-K44A was generous gifts from Pietro De Camilli. pcDNA3-HA-Arf1 (Addgene plasmid # 10830) and pcDNA3-HA-Arf1-ActQ71L (Addgene plasmid # 10832) were generated by Thomas Roberts. pGEX-5x-1-VHS GAT-GGA3 was a generous gift from Juan S. Bonifacino. pEGFP-PXN-WT was described previously (Bjorkblom et al., 2005).

### Plasmid construction

EGFP-C1tagged GIT1 phosphorylation site mutants S371A, S692A, S371A/S692A (GIT1-AA), were prepared by insertional overlapping PCR using mutagenic and flanking primers. The following flanking primers were used: forward 5**^’^** attttattgaattcaatgtcccgaaaggggccg 3**^’^** and reverse 5**^’^** attatattggtacctcactgcttcttctctcggg 3**^’^**. The following mutagenic primers were used S371A: forward 5**^’^**cagggcaagagcctgagcgcacccacagacaacctcgag 3**^’^** and reverse 5**^’^** ctcgaggttgtctgtgggtgcgctcaggctcttgccctg 3**^’^.** For S371D, primers were: forward 5**^’^** cagggcaagagcctgagcgatcccacagacaacctcgag 3**^’^** and reverse 5**^’^** ctcgaggttgtctgtgggatcgctcaggctcttgccctg 3’ S692A: forward 5’ gctgtgaccgagatggccgcactcttcccaaagaggccag 3’ and reverse 5’ ctggcctctttgggaagagtgcggccatctcggtcacagc 3’. All PCR reactions were performed using Kapa HiFi Taq Polymerase (Kapa Biosystems) and a Mastercycler Ep Gradient Auto thermal cycler (Eppendorf). Constructs were confirmed by DNA sequencing. PCR products were inserted into the EGFP-C1 vector (Clontech) using EcoRI and KpnI restriction sites. pGEX-GIT1-WT and pGEX-GIT1–S371A and S692A and double mutant were cloned in the pGEX6p-3 vector (Amersham Biosciences). For this, GIT1 PCR products were digested with EcoRI and XmaI and the vector was digested with PstI-EcoRI or PstI-XmaI and corresponding fragments were used to ligate with GIT1 sequence. Paxillin-S178A was generated by insertional overlapping PCR, using pEGFP-PXN-WT as a template. Flanking primers were:forward attttattgaattcaatggacgacctcctcgatgcc 3**^’^** and reverse 5**^’^** attatattggtaccctagcagaagaagagcttcacga 3**^’^.** Mutagenic primers for S178A were: forward 5**^’^** ccttgcctggagccctggcacccctttatggcatcccag 3**^’^** and reverse 5**^’^** ctgggatgccataaaggggtgccagggctccaggcaagg 3’. PCR products, PXN-WT and PXN-S178A were inserted into the Flag-pCMV2 vector using EcoRI and KpnI restrictions enzymes.

pAAV-CAG-Voltron was generated using pAAV-CAG-mRuby2 (Chan et al., 2017) (Viviana Gradinaru, Addgene plasmid # 99123) and pCAG-Voltron (Abdelfattah et al., 2019) (Eric Schreiter, Addgene plasmid #119033). Voltron was PCR’d out using primers forward 5’ aggtaccgccaccatggctgacgtg 3’ and reverse 5’ aaagctttcattacacctcgttctcgtagc 3’ with PCR (Phusion Hot Start II (Thermo Scientific) and T100 thermal cycler (Bio-Rad)), and used to replace mRuby2 using HindIII and Kpn1 (Fast Digest, Thermo Scientific) to produce pAAV-CAG-Voltron.

### Antibodies

For immunoblotting dilutions were as follows: 1:1000 paxillin (Bioscience), 1:500 GABAAR-β3 (Neuromab), 1:2000 GIT1 (Santa-Cruz), 1:1000 β-PIX (Bioscience), 1:1000 14-3-3ζ (Santa-Cruz), 1:2000 FLAG (Sigma Aldrich), 1:10,000 EGFP (JL-8, Clontech), 1:1000 JNK1 (Bioscience), 1:50,000 anti-rabbit and anti-mouse secondary HRP antibodies (Millipore). For immunostaining primary antibodies 1:500 GABRB3 (Neuromab), 1:500 VGAT (Millipore # AB2257) and 1:3000 VGLUT1 (Synaptic systems # 135304) were diluted in 1% FBS. GIT1, β-PIX and paxillin were diluted as described for the immunoblotting.

### Mass spectrometry

GIT1 was identified in a screen for JNK1 substrates in brain essentially in the same way as described earlier for MAP2 (Bjorkblom et al., 2005). Similarly, GIT1 phosphorylation in brains from WT and *Jnk1-/-* and DJNKI-1 infused mice was measured from TiO2-enriched phosphopeptides and analysed using an LTQ-Orbitrap XL (Thermofisher) as previously described (Padzik et al., 2016). The MS/MS spectra were exported for protein identification with Mascot version 2.3 and the UniProt database (UniProt, RRID: nif-0000-00377). Cysteine carbamidomethylation was set as a fixed modification with methionine oxidation and acetylation of the protein N-terminus added as variables. Phosphorylation of serine, tyrosine and threonine was set as variable modification when searching spectra from TiO2 enriched fractions.

### Purification of recombinant proteins in E.coli

Plasmids were transformed into E. coli Rosetta (DE3) pLysS Competent cells (Novagen-Millipore) and grown in 10 ml of Luria-Bertani (LB) broth supplemented with ampicillin. Cells were grown at 37 °C in a shaking incubator (250 rpm) overnight. The following day, 1 x l of LB was inoculated with the overnight culture. Cells were grown until OD at 590 nm was between 0.5 and 0.8 following which they were induced with 0.5 mM IPTG and harvested after 4 h by centrifugation at 4000 rpm, 4° C for 10 min. Bacteria were collected and pellets lysed in *bacterial lysis buffer* (Triethanolamine 20mM, Tris 10mM pH:7.8, NaCl 60mM, DTT 2mM, EDTA 1mM, Benzamadine 4mM and 0.05% Triton-X100) supplemented with protease inhibitors (PMSF 50 µg/ml, leupeptin, pepstatin and aprotinin at 0.2 µg/ml).

Lysozyme (2 mg/ml), DNAse-1 (50 µg/ml) and MgCl_2_ 2.5 mM were added. Cell lysates were centrifuged at 10,000 rpm, 4°C for 30 min. Resulting supernatant was incubated with 500 μl hexylglutathione agarose beads (GE Healthcare) and rotated end over end overnight at 4° C. The next day, beads were washed 5x times with *bacterial lysis buffer* at 2000 rpm, 4°C for 3 min.

Elution was performed using 1 ml glutathione 25 mM (GSH) with rotation at room temperature for 30 min. Eluted proteins were dialysed overnight in *dialysis buffer* (Tris 25 mM pH:7.4, EGTA 5 mM, DTT 2 mM, 0.1 % Triton-X100, 50 % glycerol). Protein concentration was determined from Coomassie Blue stain using bovine serum albumin for a standard curve.

### In vitro phosphorylation assay

GIT1 phosphorylation by JNK1 was identified from an *in vitro* phosphorylation assay where active GST-JNK1α1 was used to phosphorylate heat-inactivated whole brain lysate in order to identify direct targets for JNK1, as shown in Fig. 1a. This method was previously described (Bjorkblom et al., 2005; Deshpande et al., 2020). GST-GIT1 variants were phosphorylated *in vitro* using GST-JNK1α1 as previously described (Komulainen et al., 2014). Briefly, the reaction took place in reaction buffer (20 mM MOPs 2 mM EGTA, 2 mM DTT, 0.1% Triton-X-100, 25 mM ATP, 20 mM MgCl_2_ and 5 µCi ^32^P ATP) for 30 min at 30°C with 3 s, 400 rpm rotation every 1-2 min. The reaction was stopped by adding 4x *Laemmli* buffer followed by boiling for 5 min. Proteins were separated by SDS-PAGE gel and bands visualised by staining with Coomassie brilliant blue. Densitometry was carried out using a Fuji Phosphoimager.

**Figure 1.**
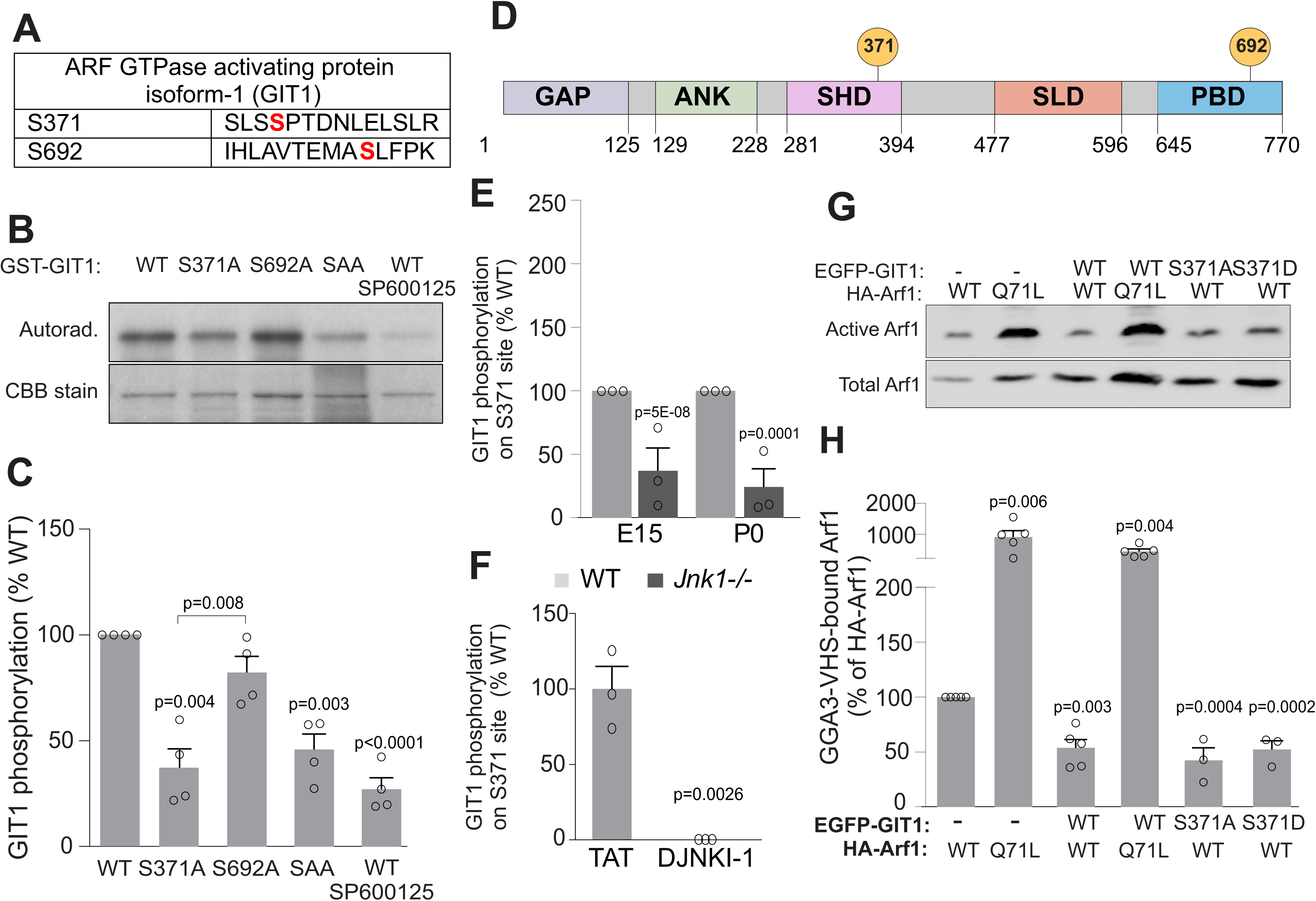
JNK1 phosphorylates GIT1 on S371 but does not alter its GAP activity Α. *In vitro* phosphorylation of brain homogenate by recombinant active JNK1 revealed phosphorylation of GIT1 on S371 and S692. Mass spectrometry-identified sequences are shown for GIT1 (accession NP_001078923.1). **Β.** Representative autoradiograph and Coomassie Brilliant Blue (CBB) stained gel image are shown for recombinant GST-GIT1-wild-type (WT), -S371A, -S692A or S371A/S692A (SAA) variants after phosphorylation by GST-JNK1. JNK1 phosphorylated GIT1-WT and GIT1–S692A but not GIT1-S371A, indicating that S371 is the preferred JNK1 phosphorylation site. Phosphorylation of GIT1 by JNK1 was prevented by JNK inhibitor 10 µM SP600125. **C.** Quantitative data from 4 repeats of experiment shown in B**. D.** Domain map of human GIT1 (UniProt identifier: Q9Y2X7-3) shows predicted JNK1 phosphorylation sites (yellow circles). Abbreviations: Arf GTPase activating protein domain (GAP), 3x Ankyrin repeats (ANK), Spa-Homology domain (SHD), Synapse Localization Domain (SLD) and Paxillin Binding Domain (PBD). **E.** Quantitative LC-MS/MS data of GIT1-S371 phosphorylation in wild-type (WT) and *Jnk1-/-* whole brains. **F.** Quantitative LC-MS/MS data of GIT1-S371 phosphorylation in brains from DJNKI-1-infused adult mice. **G.** Active Arf1 was detected from pull downs using GST-GAT-GGA3 immobilized on glutathione beads. Cells expressed GIT1-variants together with Arf1-WT or constitutively active Arf1-Q71L. **H.** Quantitative data from 3 repeats of experiment described in G. Phosphorylation-site mutants of GIT1 did not alter Arf1 activity. Error bars represent standard error of the mean (S.E.M.) and P-values were determined using Student’s t test.

### Pharmacological treatments

Neurons were treated with TAT or DJNKI-1 (Gene Cust, Laboratoire de Biotechnologie du Luxembourg, Dudelange, and Luxemburg) (20 µM) for 8h, and/or R18 (10 μM) (Enzo Life Sciences) for 24 h prior to TAT/DJNKI-1. DMSO or SP600125 (10 μM) was used overnight (-21h) (MERCK Sigma Aldrich Solutions, Darmstadt). GABA (200 µM) (Sigma)

### Primary neuron cultures

New-born Sprague Dawley rats, or *Jnk1-/- (*B6.129S1-*Mapk8^tm1Flv^*/J) *or* wild type (WT) mice (C57B6/J), of unknown sex were used at postnatal day 0. Hippocampi and cortices were dissected and mechanically dissociated after treatment with papain (Worthington) (100 U) followed by trypsin inhibitor (10 mg/ml) (Sigma). Cells were cultured onto surfaces coated with poly-D-Lysine (50 µg/ml). Hippocampal neurons that were used for staining experiments were grown in 1ml Neurobasal A medium (Gibco) supplemented with B27 (Gibco), 2 mM L-Glutamine (Biowest), penicillin (50 U/ml)/streptomycin (50 µg/ml) at a density of 100.000 cells per well in a 24-well plate. 250 ml freshly prepared medium was exchanged with culture media every four days. Cortical neurons that were used for immunoprecipitation experiments were grown in MEM medium supplemented with 10 % fetal bovine serum, 30 mM glucose, 2 mM glutamine and penicillin (50 U/ml)/streptomycin (50 µg/ml) with a density of 2.5 x 10^6^ cells per well in a 6-well plate. After 48 h, 2.5 µM cytosine arabinofuranoside (Sigma, St. Louis, MO) was added. Cells were cultured in a humidified 5 % CO_2_ incubator at 37°C. Transfections were performed at 7 days *in vitro* (DIV) using Lipofectamine^TM^ 2000 (ThermoFisher Scientific) according to the manufacturer’s instructions. DNA constructs were expressed at 30%. Neuronal experiments were performed at 16 DIV.

### Cell surface expression by biotinylation

At 16 days *in vitro*, media was removed and the cells were rinsed 2x with cold PBS supplemented with 0.5 mM MgCl_2_ and 1 mM CaCl_2_. Cells were mildly agitated for 30 min at 4°C with 1 mg/ml cell-impermeable biotin (Pierce™ Cell Surface Protein Isolation Kit, 89881). The biotinylation reaction was quenched by 5 x washes with 50 mM glycine and 0.5 % BSA dissolved in supplemented PBS followed by 3x PBS washes. We lysed cells in radioimmunoprecipitation assay (RIPA) low salt buffer (50 mM Tris, pH7.4, 1 mM EDTA, 2 mM EGTA, 150 mM NaCl, 1 % NP40, 0.5 % deoxycholate and 0.1 % SDS) supplemented with protease inhibitors: 1 µg/ml of aprotonin (Sigma Aldrich), leupeptin (Sigma Aldrich), pepstatin (Apollo) and 100 µg/ml of phenylmethylsulfonyl fluoride (Calbiochem) and incubated for 15 min at 4°C. Cell lysates were homogenized using a 27g needle and precleared by centrifugation at 13,200 x g, 4° C for 15 min. 10 % of the supernatant was used to measure total input. Biotinylated proteins were isolated from supernatants by incubating with NeutrAvidin Agarose beads (3:1, cell lysate: beads ratio) for 2 h at 4° C with rotation. We washed beads (3 x 5 min) in RIPA high salt buffer (50 mM Tris pH7.4, 1 mM EDTA, 2 mM EGTA, 500 mM NaCl, 1 % NP40, 0.5 % deoxycholate and 0.1 % SDS) followed by 1 x with RIPA low salt buffer. 800 x g spins were used at 4°C. Beads were eluted by boiling in Laemmli buffer containing beta-mercaptoethanol for 6 min prior to SDS-PAGE and immunoblotting.

### Human Embryonic Kidney cell culture

Human embryonic kidney (HEK293) cells (ATCC: CRL-1573) were grown and transfected as previously described (Padzik et al., 2016).

### Immunostaining

Hippocampal neurons at 16 DIV on coverslips (VWR, 13 mm, thickness No 1.5) were fixed in pre-warmed at 37° C paraformaldehyde (4%) for 20 min at RT, except for surface staining when fixation was 6 min as previously described (Smith et al., 2014). Cells were washed 3x times with 1x Phosphate Buffer Saline (PBS) and incubated with 1% Triton-X100 in PBS for 3 min (except when cell surface staining was performed, when this step was omitted). Cells were washed 3x times with PBS and blocked with 10 % fetal bovine serum for 1 h at RT, washed 3x times with PBS and incubated overnight with primary antibody at 4° C. Primary antibodies were diluted (1:500) in 1% FBS. Next day, cells were incubated with Alexa-488, Alexa 546 and Alexa 647 at 1:1000 dilution for 1 h at RT. Cells were washed 3x times with PBS and mounted on microscope slides (VWR) with Mowiol (Calbiochem).

### Confocal imaging for spine and dendrite analysis

Confocal images of 16 day *in vitro* (D.I.V.) hippocampal neurons were obtained using a Zeiss 780 LSM with 63X oil objective (1.4 NA) (Fig. 2) or a Zeiss 880 LSM using the AiryScan mode and 63X oil objective (1.4 NA) using 488 nm (12% laser power) (EGFP), 543 nm (27% laser power) (mRuby-Lifeact) or 633 nm laser lines (11% laser power) (Alexa-647) (all other figures). Scans used 4-line averaging with a minimum pixel dwell time of 0.60 μs, pixel size of 0.043 μm x 0.043 μm and airy unit of 1. Images were taken from secondary proximal apical dendrites that were 25 μm from the soma, avoiding bifurcating dendrites or the distal tip of the dendrite. For analysis, maximum z-sections were generated from 18 x z-sections (optical section 0.173 μm each), unless otherwise indicated in figure legend. For imaging of GABAARs, z-stacks were made from 6 sections, 0.833 μm apart obtained with the settings as described above.

**Figure 2.**
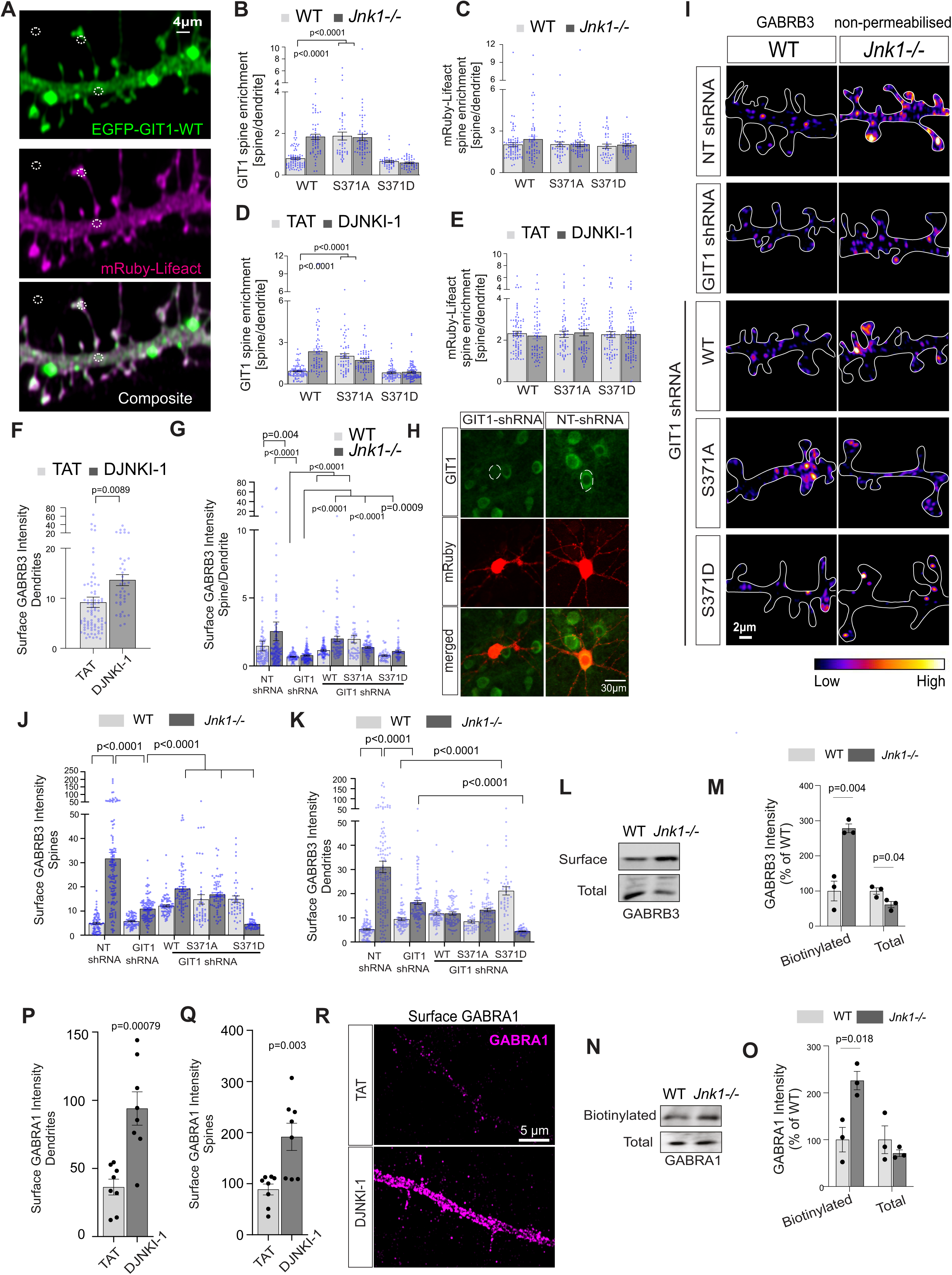
Endogenous GIT1 and GIT1-S371A enrich in dendritic spines of *Jnk1-/-* or JNK-inhibitor treated neurons. **A.** Representative maximum projection images (25x optical slices, z=0.173 µm) depict EGFP-GIT1-WT and mRub-Lifeact in 16-DIV hippocampal neurons. Regions of interest (R.O.I.s; circles) depict where in the dendrite, spine and background mean intensity signals were acquired. **B, D.** Relative enrichment of EGFP-GIT1 variants in spines or dendrites, was calculated as a ratio of the mean intensity from the spine R.O.I. to the mean intensity from the R.O.I in the dendrite, after background subtraction, for the given treatments *Jnk1-/-* or TAT/DJNKI-1 (20 µM), 8 hours. **C, E.** Relative enrichment of mRuby-Lifeact fluorescence in spines was measured as for B and E. Mean data +/- S.E.M. from a total of ∼80 spines from several neurons per condition are shown. Neurons originated from at least two independent mouse litters. P-values were calculated from Student’s two-tailed t test. **F, G.** We next checked the overall effect of DJNKI-1 inhibitor or *Jnk1* knockout on β3 GABAAR (GABRB3) expression at the cell surface in the dendritic compartment. Both conditions increased general surface expression of GABRB3. Gene names are used for GABAAR S.U.s in the figures to save space. **H.** In order to test the requirement for GIT1 in GABRB3 trafficking, we characterised shRNA targeting GIT1. **I.** Representative maximum projection images of GABRB3 staining from 25 x optical sections (z=0.173 μm) are shown for WT and *Jnk1-/-*hippocampal neurons at 16 DIV. Staining was performed in non-permeabilized cells expressing mRuby, with non-targeting (NT)-shRNA or GIT1-shRNA and EGFP-GIT1 variants, as shown. White outlines were traced from mRuby fluorescence. Surface staining of GABRB3 is shown in pseudocolour. Representative images of total GABRB3 staining from an independent set of permeabilized cells is shown for comparison. **J, K**. Quantitative data from these experiments is shown for spines and dendrites. Mean data +/- S.E.M are shown from ∼70 dendritic spines per condition, in cells from at least two independent mouse litters. **L-O.** Cell surface expression of GABRB3 or GABRA1 was measured from biotinylated samples (L, N). Representative blots show GABRB3 or GABRA1 from lysates from before streptavidin-biotin enrichment (total) or after streptavidin-biotin enrichment (surface) are shown (M, O). Mean data from 3 repeats +/- S.E.M are shown. P-values were calculated from Student’s two-tailed t test. **P-R.** (P,Q) Quantitative data GABRA1 levels in 16 DIV WT and *Jnk1-/-* hippocampal neurons in non-permeabilized cells treated with TAT/DJNKI-1 (20 µM), 8 hours. Mean data from 8 cells per condition +/- S.E.M are shown. P-values were calculated from Student’s two-tailed t test. (R) Representative maximum projection images (6x optical slices, z=0.211 µm) of GABRA1 staining are shown.

For VGAT/VGLUT1 and GABAAR-β3 co-localization analysis, confocal images of 16 day in vitro hippocampal neurons were obtained using a Zeiss 880 LSM with AiryScan detector and 63X oil objective (1.4 NA). Laser lines were 488 nm at 6.4 % power for Alexa-488 used to detect VGAT and VGLUT1; 543 nm at 15.4% power to detect mRuby2 and 633 nm with 14.7% l power with Alexa-647 to detect GABAAR-β3. For Alexa-488, beam splitter SBS-SP-615 and filter BP 420-480 + BP 495-550 were used, while for mRuby2 and Alexa-633, BP 555-620 + LP 645 and 633 BP 570-620 + LP 645 filters were used. Scans used 4-line averaging with minimum pixel dwell of 0.76 μs, pixel size of 0.04 μm x 0.04 μm. As above, images were taken from secondary proximal apical dendrites that were 25μm from the soma, avoiding bifurcating dendrites or the distal tip of the dendrite. Images were captured in frame mode by acquiring Alexa-647, mRuby2 and Alexa-488 channels sequentially. For analysis, maximum z-projections were generated from 6 x 0.211 µm z-sections. The experimenter was blinded to treatments before imaging and until the analysis was completed.

### Widefield analysis with Voltron

Neurons were transduced with AAVs expressing Voltron from the plasmid pAAV-CAG-Voltron. For imaging, cells were incubated with 100 nM of Janelia Fluor dye (JF549) with HaloTag ligand (Promega) for 30 min prior to imaging at 37°C. Media was changed 10 min before imaging to 145 mM NaCl, 2.5 mM KCl, 10 mM glucose, 20 mM HEPES, pH 7.4, 2 mM CaCl_2_, 1 mM MgCl_2_. Hanks Balanced Salt Solution (GIBCO, ThermoFisher). Cells were imaged at 15 or 17 DIV using a Nikon Eclipse Ti2-E microscope with a 40x oil objective and acquired with a Hamamatsu sCMOS Orca camera. Lumencor Spectra X LED was used with excitation filters 550/15 nm, and emission filters 595/40 nm were used. Fluorescence over time was measured at 60 ms intervals from within regions of interest (R.O.I.) in the dendrite or at the soma. An equal-sized R.O.I. from outside the cell was acquired as background.

### Image Analysis

Fluorescence intensity in dendritic spines and in proximal dendritic shafts was measured from maximum z-projection images using ImageJ (NIH) with the circle tool to define R.O.I.s between 0.4 to 0.7 µm^2^. R.O.I.s were drawn for background, spine-head and adjacent dendrite regions as shown in figure 2A. Mean background R.O.I. fluorescence intensity was subtracted from spine and dendrite mean intensity measurements in equal sized R.O.I.s. Mean fluorescence intensity in dendritic spine R.O.I. was divided by mean fluorescence intensity in R.O.I. in the adjacent dendrite to provide spine enrichment values as shown in Fig. 2. The same procedure was applied to measure spine enrichment of mRuby-Lifeact. and to analyse relative fluorescence intensity of endogenous GIT1, paxillin, 14-3-3ζ and β-PIX in figures 3, 4, 5 and 7. For GABAAR receptor analysis however (Fig.7), mean fluorescence intensity was analysed from both permeabilized cells (representing total labelling) and non-permeabilized cells (representing surface labelling). Dendritic morphology was scored manually using Neurolucida 8.03 (MBF Bioscience, Williston, VT USA). For dendritic spine morphology analysis, spines were categorised into three morphological categories: mushroom, thin and stubby as previously described (Peters and Kaiserman-Abramof, 1970; Hering and Sheng, 2001), where mushroom spines had a head:neck ratio > 2; thin spines had a head:neck ratio < 2. Spines with no neck that were attached to the dendritic shaft were defined as stubby. Protrusions that were too wide or flat were excluded. Between 40 and 80 spines were measured per condition as described in the legends. Spine measurements were pooled from at least 5 neurons from 4 different coverslips.

**Figure 3.**
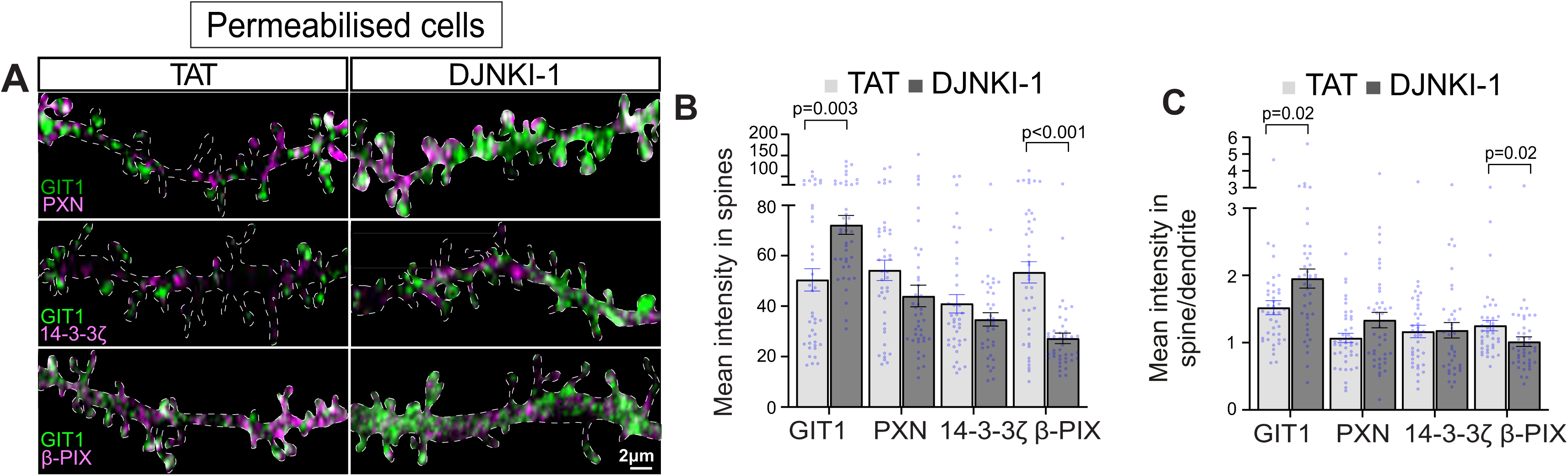
JNK inhibition enriches endogenous GIT1 in spines where β-PIX is reduced. **A.** Representative maximum projection images (32 x optical sections of 0.173 μm) of neurons expressing mRuby2 to visualize morphology. Neurons at 16 DIV were immunostained for endogenous GIT1 (green) and paxillin (PXN), 14-3-3ζ or β-PIX (magenta). **B.** Quantitative data show mean intensities of GIT1, PXN, 14-3-3ζ and β-PIX in spines. DJNKI-1 increases GIT1 in dendritic spines compared to control (TAT) and reduces β-PIX in the same cell compartment. **C.** Mean intensity ratios of GIT1, PXN, 14-3-3 ζ and β-PIX in dendritic spine/shaft. Mean data +/- S.E.M.s from ∼42 spines per condition.

**Figure 4.**
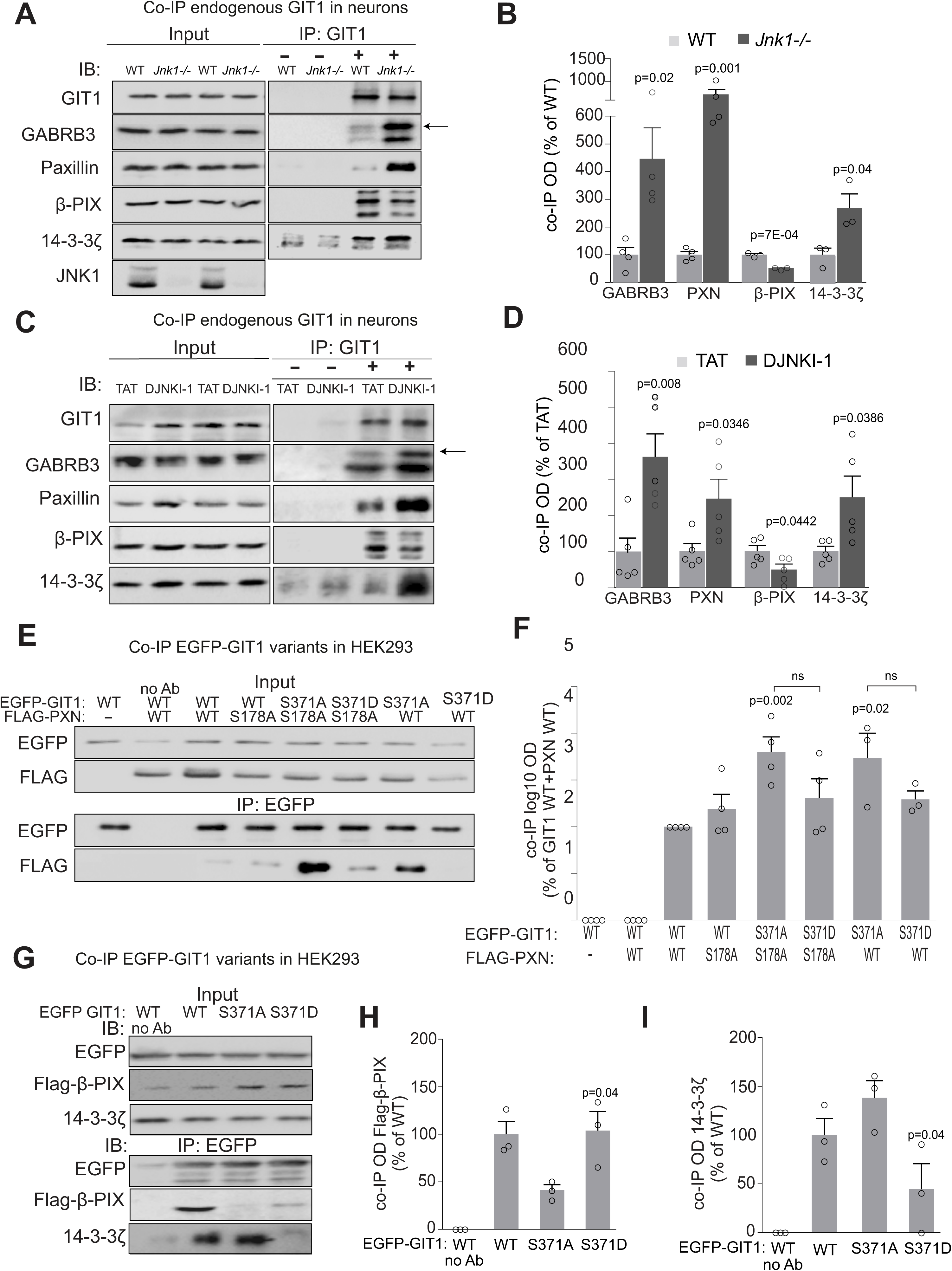
Phosphorylation of GIT1 by JNK1 decreases its association with molecular protein partners: GABAAR-β3, PXN and 14-3-3ζ Α. We examined whether GABRB3, paxillin, β-PIX or 14-3-3ζ co-immunoprecipitated with GIT1 using cortex-derived homogenates from WT and *Jnk1-/-* adult mice. +/- denotes addition or not of antibody. Representative immunoblots of input and GIT1 immunoprecipitates probed with indicated antibodies are shown. **B.** Quantitative data show the extent of GABRB3, PXN, β-PIX and 14-3-3ζ co-purification with GIT1 from brain. For 14-3-3ζ quantification, the background bands present in samples without GIT1 antibody were subtracted from bands in samples with GIT1 antibody. There was increased interaction between GIT1 and GABRB3, PXN and 14-3-3ζ in *Jnk1-/-* brain compared to WT. In contrast, interaction between GIT1 and β-PIX decreased. Histogram bars represent mean data from 4 experimental repeats +/- S.E.M.. **C.** The effect of TAT or DJNKI-1 on GIT1 interaction with binding partners was determined. Cortical neurons at 16 DIV were treated with 20 µM TAT or DJNKI-1 for 8 hours. Representative immunoblots of GIT1 co-precipitations are shown. **D.** Band quantification from blots in Fig. 6C. DJNKI-1-treated neurons show increased interaction between GIT1 and GABRB3, paxillin and 14-3-3ζ, whereas β-PIX interaction with GIT1 reduced. Bars show mean data from 5 repeats ± S.E.M.. **E.** We tested whether GIT1-S371 phosphorylation site mutants altered these interactions. HEK-293 cells were transfected with EGFP-GIT1-WT, EGFP-GIT1-S371A or EGFP-GIT1-S371D and either Flag (as a control), Flag-PXN-WT or Flag-PXN-S178A, as indicated. Representative blots of input and co-immunoprecipitations are shown. **F.** Quantitative band intensities from E are shown using log10 scale. PXN-S178A interacted most highly with GIT1-S371A. Mean data ± S.E.M. from 4 repeats are shown. Student’s t-test p-values above bars are compared to samples with EGFP-GTI1-WT and Flag-PXN-WT. **G.** The same approach was used to test endogenous 14-3-3ζ and Flag- β-PIX interaction with GIT1. Representative blots are shown. **H.** Quantitative data for Flag- β PIX interaction with EGFP-GIT1 variants. **I.** Quantitative data for 14-3-3ζ interaction with EGFP-GIT1 variants. Endogenous 14-3-3ζ interacts most highly with EGFP-GIT1-S371A. Bars ± S.E.M. represent means from 3 repeats. p-values were calculated from Student’s two-tailed t-test and are compared to conditions where WT is expressed. IB: immunoblot, IP: immunoprecipitation.

**Figure 5.**
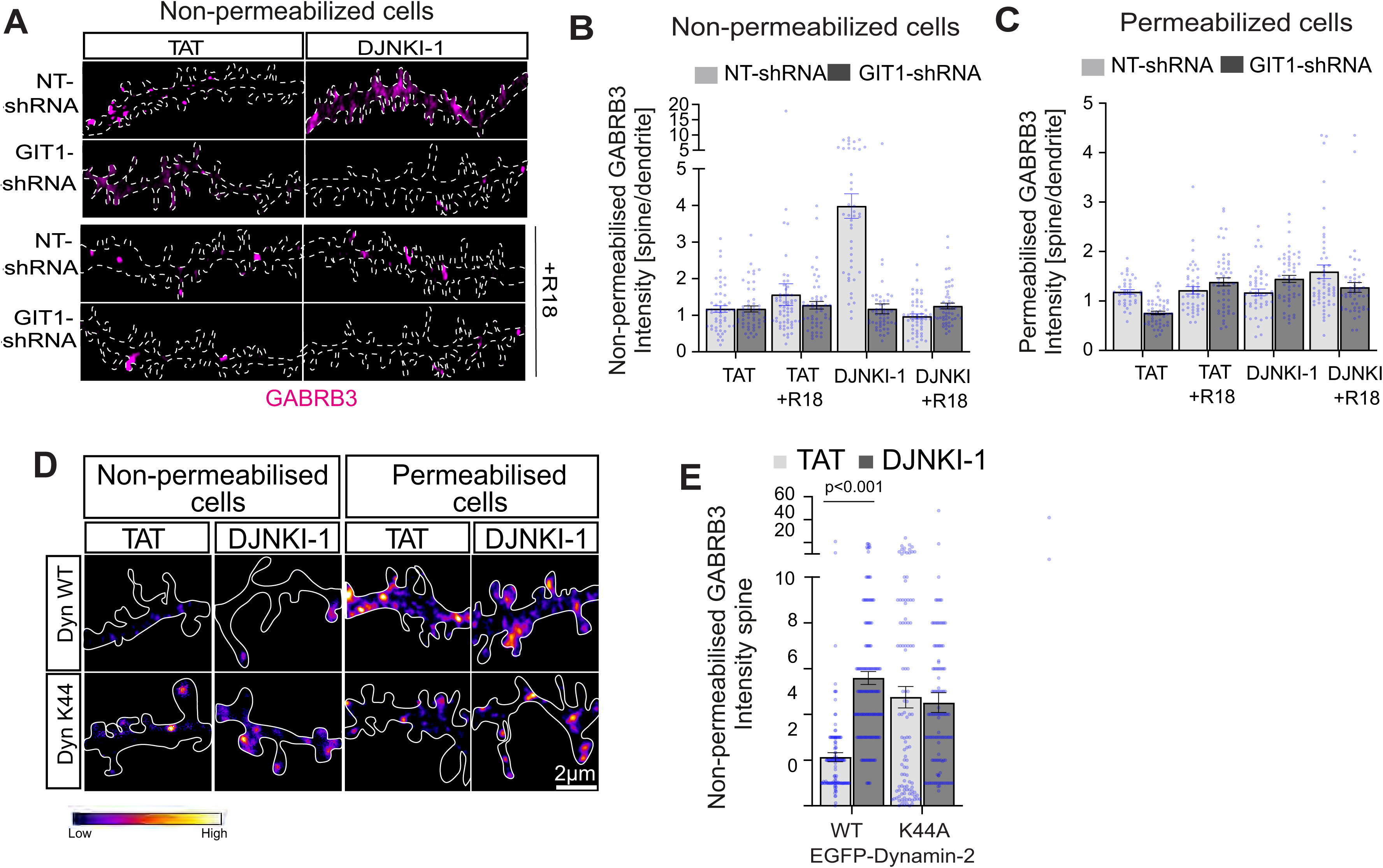
R18 inhibitor of 14-3-3ζ client binding prevents GABAAR-β3 surface enrichment in JNK inhibitor treated neurons. **A.** Representative maximum projection images (22 x optical sections of 0.173 μm) showing staining of GABRB3 in non-permeabilised 16-DIV hippocampal neurons (magenta). Cells co-express mRuby2 with NT-shRNA or GIT1shRNA, as indicated. Cell tracing was based on mRuby2 images. Cells were treated with 10 µM R18 for 24 h and 20 µM TAT or DJNKI-1 for 8 h. **B.** Quantification of GABRB3 enrichment in dendritic spines. R18 prevented DJNKI-1-induced GABRB3 increase at the dendritic spines. **C.** R18 prevented GIT1-S371A-induced increase in GABRB3 at the cell surface. Histogram bars represent means from ∼51 spines per condition from several neurons. P-values were calculated from Student’s two-tailed t-tests were compared to control (TAT) unless otherwise indicated. **D.** Neurons were co-transfected at 7 DIV with EGFP-Dynamin 2-WT or EGFP-Dynamin 2-K44A and treated with 20 µM TAT or DJNKI-1 for 8 h. Representative images are shown. **E.** Mean intensities for GABRB3 are shown from non-permeabilised cells in mushroom spines. GABRB3 surface expression increased in DJNKI-1-treated neurons, even when the K44A mutant is expressed. Mean data +/- S.E.M.s from ∼70 spines per condition are shown. p-values (calculated from Student’s two-tailed t test) were compared to control (TAT) unless otherwise indicated by brackets.

### Immunoprecipitation

Cortex from adult wild-type or *Jnk1-/-* C56BL6/N mice was frozen in liquid nitrogen after extraction and stored at -80 °C until use. Tissues were homogenised in *hypotonic lysis buffer* (20 mM Hepes pH:7.4, 2 mM EGTA, 50 mM β-glycerophosphate, 1 mM DTT, 1 mM Na_3_VO_4_, 1 % Triton-X-100, 10 % glycerol, 50 mM NaF. 1 mM benzamidine) supplemented with protease inhibitors: aprotinin (Sigma Aldrich), leupeptin (Sigma Aldrich), pepstatin (Apollo) at 1 μg/ml and phenylmethylsulfonyl fluoride (PMSF, Calbiochem) at 100 μg/ml for 15 min at 4°C. Homogenization was performed using 3x 30 s pulses with an Ultra Turrax tissue homogeniser. Supernatant was precleared by centrifuging at 13,500 x g for 15 min at 4° C. Protein concentration was quantified using the Bio-Rad Protein Assay Dye Reagent Concentrate (Bio-Rad). 10 % of the supernatant was kept to assess “total input” by immunoblotting. EGFP-GIT1 variants were immunoprecipated from precleared lysates using 0.5 μg EGFP antibody per sample. For immunoprecipitation from tissues and neuronal cultures, GIT1 was immunoprecipitated with 0.5 μg GIT1 antibody. After antibody addition, samples were rotating end over end overnight at 4° C. The following day, lysates were incubated for 2 h with 10 μl bed volume of protein A-agarose (Upstate Biotechnology) at 4°C. Complexes were washed with PBS 5x times for 5 min each and collected using 800 x g at 4°C, followed by boiling for 5 min in 50 μl 1 x *Laemmli* buffer prior to SDS-PAGE and western blotting.

### Arf1 activity

Was measured using GST-VHS–GAT-GGA3 as previously described (Dell’Angelica et al., 2000). Briefly, VHS-GAT-GGA3 was produced as described above and immobilised on glutathione sepharose beads (GE Healthcare). HEK293 cells on 10 cm dishes were transfected with pcDNA3-HA-Arf1-ActQ71L or pcDNA3-HA-Arf1 making up 30% of the DNA mix, with either EGFP-N1 empty vector or EGFP-N1-GIT1-WT-S371A or -S371D (8% of DNA mix). After expression, cells were homogenized in 300 μl of *hypotonic lysis buffer*, supplemented with protease inhibitors. 250 μl of supernatant was incubated with 50 mg of GST-VHS–GAT-GGA3 glutathione beads for 30 min on a rotating rack at 4°C. Pellets were centrifuged, 800 x g at 4°C, following which 2x washes were performed with *lysis buffer*. Pellets were resuspended in 50 μl 1x Laemmli buffer and boiled for 5 min prior to SDS-PAGE and western blotting.

### Immunoblotting

For western blotting, lysates underwent SDS-PAGE followed by transfer to polyvinylidene fluoride membrane (Bio-Rad). Membranes were incubated with 1 x PBST (Phosphate Buffered Saline Tween-20) containing 5 % skimmed milk, for 1 h at RT. Primary antibody incubation was performed at 4 °C overnight. The following day, membranes were washed with 1 x PBST, and incubated with horse radish peroxidase-labelled goat anti-mouse or rabbit IgG (Millipore) (1:50.000) for 1 h at RT according to the manufacturer’s instructions. Bands were exposed to SuperSignal West Femto Maximum Sensitivity Substrate (ThermoFisher Scientific) and visualised using the ChemiDoc^TM^ MP Imaging System (Bio-rad). Immunoblots were quantified by densitometry using ImageJ Fiji 1.51s (NIH) software.

### Electrophysiology

Electrophysiological experiments were done using the whole-cell patch-clamp technique. Cultures of 14 – 16 DIV neurons were transferred from the growing medium to pre-warmed 33°C artificial cerebrospinal fluid (ACSF), containing: 126 mM NaCl, 1.6 mM KCl, 1.2 mM MgCl_2_, 1.2 mM NaH_2_PO_4_, 18 mM NaHCO_3_, 2.5 mM CaCl_2_ and 11 mM D-glucose (Heikkinen et al., 2009). Cells were visualized with an upright brightfield microscope (BX51WI, Olympus, Tokyo, Japan) and a digital camera (Hamamatsu C8484, Hamamatsu City, Japan). Whole-cell voltage-clamp registrations were made using 3-5 MΩ borosilicate glass electrodes filled with intracellular solution (IS). All recordings were done with a Multiclamp 700 B amplifier and Digidata 1440A (both Axon Instruments, Molecular Devices, San Jose, USA), digitized with 10 kHz sampling rate using pClamp 10 software. Access resistances were monitored throughout the experiment, and the recording was discarded if it changed >20% during the experiment.

For tonic current experiments IS contained: 140 mM CsCl, 4 mM NaCl, 0.5 mM CaCl_2_, 10 mM HEPES, 5 mM EGTA, 2 mM Mg-ATP, pH 7.2–7.25 adjusted by CsOH, osmolarity 280 mOsm. After achieving whole-cell configuration, cells were clamped at -70 mV. To block all glutamatergic currents, kynurenic acid (2 mM) was added to the constantly perfused ASCF, together with tetrodotoxin (1 µM) to abolish all synaptic activity, NNC-711 (10 µM) to block GABA reuptake and GABA at a low concentration (500 nM) to activate extrasynaptic GABAARs. After recording of the baseline inward current (3-4 min), gabazine (250 µM) was added into the ACSF. Tonic current was measured as described previously (Bright & Smart, 2013). Shortly, we generated in Clampfit software (pClamp 10, Molecular Devices) an all-points amplitude histogram before (30-s epoch) and after gabazine action (10-s epoch) and fitted them separately with Gaussian. The mean values from the Gaussian fits were then used to define the current amplitudes in baseline and gabazine conditions. Tonic current was calculated according to the formula: **I_tonic_= I_baseline_– I_gabazine_**. Comparison of the two genotypes was done using unpaired two-tailed *t*-test, after removal of two outliers from the WT group using ROUT method (Q=1%) in Prism 8.1.0 (GraphPad Software, San Diego, USA). All results are shown as mean ± standard error of the mean (S.E.M.).

For sIPSCs experiments IS composition was as follows: 130 mM K-gluconate, 6 mM NaCl, 10 mM HEPES, 0.5 mM EGTA, 4 mM Na_2_-ATP, 0.35 mM Na-GTP, 8 mM Na_2_-phosphocreatine (pH adjusted to 7.2 with KOH, osmolarity ∼285 mOsm). At these conditions, calculated reversal potential for Cl^-^ was -84 mV, making the IPSCs to be outward currents at −40 mV holding potential. Recordings were started 15 min after transferring cultures from the culture medium to 33°C ACSF to ensure stabilized environment. To measure amplitudes of spontaneous events and inter-event intervals 10-min traces were analyzed. To analyze kinetics, we randomly chose representative IPSCs (20% of all events) for each cell and further analyzed rise and decay times as well as half-width of each event. Primary analysis was done in Mini Analysis 6.0.3 software (Synaptosoft Inc., Decatur, GA, USA) with further statistical analysis in Prism 8.1.0.

### Statistics

Statistical differences were evaluated using two-tailed Student’s t test or Wilcoxon test were used as indicated in the legends. P-values < 0.05 were taken as a significance threshold. Data was expressed as the mean ± S.E.M.. Actual p-values comparing treatment to control condition are depicted on the histograms above the “treatment” unless otherwise indicated by brackets.

## RESULTS

### JNK1 phosphorylates GIT1 on S371 *in vivo* in mouse brain

While carrying out an *in vitro* screen for JNK targets in mouse brain, we identified GIT1 (UniProt Q5F258) as a candidate substrate (Fig. 1). Specifically, we found that recombinant active GST-JNK1 phosphorylated peptides (SLS**S**PTDNLELSAR and IHLAVTEMA**S**LFPK) on S371 and S692 *in vitro* in a brain lysate (Fig. 1A). To validate that JNK1 directly phosphorylated GIT1 on these sites, we purified recombinant GST-tagged GIT1-WT, GIT1-S371A, GIT1-S692A and GIT1-S371A/S692A, and phosphorylated them *in vitro* with active JNK1 in a reconstituted assay (Fig. 1B). JNK1 phosphorylated GIT1-WT and GST-GIT1-S692A, but phosphorylation of recombinant GIT1-S371A was greatly reduced (Fig. 1C), indicating that GIT1-S371, which lies within the Spa2 homology domain (SHD), is a preferred JNK1 phosphorylation site, whereas S692 is not (Fig. 1D). To further evaluate this finding, we examined GIT1 phosphorylation in wild type and *Jnk1-/-* mouse brain. GIT1 phosphorylation on S371 was reduced in brains from mice lacking *Jnk1* at E15 and at P0 (Fig. 1E). Finally, we measured GIT1 phosphorylation from brains of adult mice that had been infused with a pan-JNK peptide inhibitor (DJNKI-1). GIT1-S371 phosphorylation was completely ablated in the cortex from these mice (Fig. 1F). Overall, these data indicate that JNK1, a highly active JNK isoform in brain (Coffey et al., 2000; Brecht et al., 2005; Muraleva et al., 2025), phosphorylates GIT1 on S371.

### GIT1 phosphorylation on S371 does not alter its GAP activity

GIT1 is an Arf GTPase activating protein (GAP) that can inactivate Arf1 and Arf6 (de Curtis and Paris, 2005). We therefore tested whether JNK1 phosphorylation of GIT1 on S371 may alter its GAP activity. Arf1 activity was assessed by measuring Arf1 binding to the ADP-ribosylation factor binding protein (GGA3-VHS) which binds specifically to active Arf1 (Dell’Angelica et al., 2000). As a positive control, constitutively active EGFP-Arf1^Q71L^ showed increased GGA3-VHS binding (Fig. 1G, H). GIT1-WT inhibited Arf1 binding to GGA3-VHS by 50% as expected, however this was independent of S371 phosphorylation (Fig. 1G, H), indicating that GIT1 phosphorylation by JNK1 does not regulate its GAP activity. This is consistent with S371 being outside of the GIT1 GAP domain (Fig. 1D).

### Genetic deletion of *Jnk1* or JNK inhibitor enriches surface GIT1 expression via a mechanism involving S371

GIT1 functions as a scaffold that translocates signalling proteins to dendritic spines (Hoefen and Berk, 2006). We therefore tested whether phosphorylation by JNK1 influenced the subcellular localisation of GIT1. Hippocampal neurons from wild type or *Jnk1-/-* mice were transfected with EGFP-GIT1-WT, -S371A or -S371D, and the enrichment of these variants in spines relative to the proximal dendrite was determined. In *Jnk1-/-* neurons, EGFP-GIT1-WT was enriched by twofold in the spine-head relative to the proximal dendrite (Fig. 2A-B, Supplementary figure 1). Mutation of EGFP-GIT1-S371→A was sufficient to drive it to spines in wild type neurons, unlike GIT1-S371D (Fig. 2B, Supplementary figure 1). As a control, we measured mRuby-Lifeact which binds to F-actin. There was no enrichment of this protein in spines in the presence of GIT1-371 variants (Fig. 2C, Supplementary figure 1). These results suggest that JNK1 may by phosphorylating GIT1, drive it out from spines to the dendritic shaft.

We next examined whether a brief pharmacological inhibition of JNK would affect GIT1 localisation. Hippocampal neurons treated with a peptide inhibitor of JNK (D-JNKI-1) showed a two-fold accumulation of EGFP-GIT1 in spines, whereas EGFP-GIT1-S371D localisation did not change (Fig. 2D). mRuby-Lifeact distribution also remained unchanged (Fig. 2E). This data indicates that JNK1 phosphorylation of GIT1 on S371 reduces its presence in spines.

### JNK1 activity attenuates β3 and α1- containing GABAAR levels at the cell surface

We wanted to examine if surface expression of the GABAAR was altered by these treatments as GIT1 was previously associated with GABAAR trafficking (Smith et al., 2014). We used two independent approaches to measure surface expression. First, we analysed GABAAR-β3 staining in non-permeabilised hippocampal neurons with an antibody recognising the extracellular domain of the widely expressed GABAAR-β3 subunit (Olsen and Sieghart, 2008). We analysed GABAAR-β3 surface expression in dendrites and spines, as we had observed GIT1 motility between these compartments in live imaging analysis, and additionally because 18% of inhibitory synapses are on spines (Knott et al., 2002; Kwon et al., 2019). We found that either inhibition or genetic deletion of *Jnk1* increased surface GABAAR-β3 levels in dendrites and spines (Fig. 2F, G). GIT1 knockdown attenuated the upregulation of surface β3-containing GABAARs in spines and dendrites, in *Jnk1-/-*cells (Fig. 2G-I; Supplementary Fig. 1), and this was partly rescued by expression of EGFP-GIT1 in spines (Fig. 2J, K).

We next measured the regulation of β3 and α1-containing GABAARs by JNK1 using surface biotinylation (Fig. 2L-O). This revealed that surface levels of GABAAR-α1 were also increased in spines and dendrites of hippocampal neurons following DJNKI-1 treatment (Fig. 2P-R). Combined, these results confirm that JNK1 activity attenuates cell surface expression of β3- and α1-containing GABAARs.

### JNK inhibition enriches endogenous GIT1 in spines and depletes β-PIX

Up to this, our experiments had only measured exogenously expressed GIT1. To determine whether endogenous GIT1 also enriched in spine-heads following JNK inhibition, we treated neurons with DJNKI-1 for 8 hours and detected endogenous GIT1 using immunofluorescence staining. As identified earlier for EGFP-GIT1 (Fig. 2), DJNKI-1 also increased levels of endogenous GIT1 in spines (Fig. 3A-C). GIT1 associates with several binding partners to fulfil its scaffolding role. Prominent among these are β-PIX, paxillin and 14-3-3ζ (Angrand et al., 2006; Zhou et al., 2016). We therefore examined whether the localisation of any of these proteins in dendrites and spines was altered by JNK. We found that endogenous GIT1 levels in spines increased upon DJNKI-1 treatment, howeve β-PIX levels were reduced (Fig. 3A-C). In contrast, expression of paxillin and 14-3-3ζ were equally distributed between spines and dendrites did not change after inhibitor treatment (Fig. 3A-C). These data show that β-PIX is the only GIT1 binding partner (from those measured) that changes distribution upon JNK1 inhibition. This may reflect reduced binding of βPIX to GIT1.

### JNK1-phosphorylated GIT1 shows altered interaction with GABAAR-β3, paxillin, β-PIX and 14-3-3ζ

We next tested whether endogenous GIT1 interacted in a complex with the GABAAR-β3-containing receptor in brain and if so, whether this interaction was controlled by JNK1. Co-immunoprecipitation analysis of GIT1 complexes from cerebral cortex showed that GABAAR-β3 co-purified with GIT1 and that this interaction increased four-fold in *Jnk1-/-* forebrain (Fig. 4A and B). Moreover, GIT1 interaction with paxillin and 14-3-3ζ also increased in *Jnk1^-/-^* forebrain, whereas interaction with β-PIX decreased (Fig. 4A, B). We also tested the effect of JNK inhibitor on GIT1 interactions using cultured cortical neurons. Once again, in DJNKI-1-treated cells, GIT1 interaction with GABAAR β3, paxillin and 14-3-3ζ increased, whereas interaction with β-PIX decreased (Fig. 4C, D). Together these data indicate that JNK1 phosphorylation of GIT1 increases its interaction with β-PIX, whereas binding to 14-3-3ζ GABAAR-β3 and paxillin increases when JNK is inhibited. The SHD domain of GIT1, which harbours S371, has been reported to mediate its interaction with paxillin (Totaro et al., 2007). We tested this by co-expressing GIT1-S371 variants in HEK-293 cells alongside Flag-tagged paxillin WT or Flag-PXN-S178A, a site that is phosphorylated by JNK1 (Huang et al., 2003) (Fig. 4E, F). This data confirmed the earlier findings. Thus EGFP-GIT1-S371D did not interact with paxillin-WT, whereas GIT1-S371A interacted strongly. This interaction increased even further in the S178A mutant (Fig. 4E, F). These data indicate once more that the JNK1 phosphorylation of GIT1-S371 blocks its interaction with paxillin.

We extended this analysis by examining the interaction between EGPF-GIT1 variants and Flag-β-PIX or 14-3-3ζ expressed in HEK-293 cells. β-PIX showed an increased interaction with GIT1-S371D compared to that with GIT1-S371A (Fig. 4G, H), whereas 14-3-3ζ interaction with GIT1-S371D was lower than with GIT1-S371A (Fig. 4G, I). Collectively, these data suggest that JNK1-dependent phosphorylation of GIT1 on S371 serves to disrupt its interaction with a complex containing GABAAR-β3, paxillin and 14-3-3ζ but promote its interaction with β-PIX, which excludes it from spines.

### 14-3-3ζ binding is required for GABAAR-β3 stabilization at the cell surface

To test whether 14-3-3ζ binding to GIT1 is required for GABAAR-β3 stabilization at the cell surface, we used the R18 peptide (PHCVPRDLSWLDLEANMCLP) which inhibits 14-3-3 client binding (Kaplan et al., 2017). R18 treatment completely abolished the accumulation of GABAAR-β3 at the cell surface of spines in DJNKI-1-treated neurons (Fig. 5A-C). These results together with the findings in Fig. 4A and B indicate that 14-3-3ζ binding to GIT1-S371A is necessary to achieve GABAAR-β3 surface stabilization by JNK1.

### JNK regulation of surface GABAAR-β3 may involve dynamin-mediated endocytosis

We wanted to know if the mechanism whereby JNK-controlled β3-containing GABAAR surface expression involved endocytosis, so we used dominant negative dynamin-2 (Dyn2-K44A) which blocks dynamin-mediated endocytosis. Dyn-2K44A expression led to accumulation of GABAAR-β3 at the spine surface as expected, and there was no additivity upon D-JNKI-1 treatment (Fig. 5D, E). These results suggest that JNK1 drives the endocytosis-based removal of β3-containing GABAAR from the surface of spines rather than a lateral flow mechanism.

### JNK1 prevents accumulation of β3-containing GABAARs at glutamatergic synapses

We next examined whether the β3-containing GABAARs at the cell surface were localised with glutamatergic or GABAergic synapse markers in dendritic spines. Co-staining with the presynaptically localised vesicular glutamate or GABA transporters (VGLUT1 and VGAT respectively), was used to identify potential synaptic localisation (Fig. 6A). In spines, the percentage of GABAAR-β3 that colocalised with VGLUT1 increased significantly in *Jnk1-/-*neurons, while there was no change in GABAAR that colocalised with VGAT (Fig. 6A-C). In the dendrite, GABAAR-β3 levels increased in the extrasynaptic sites and decreased at VGAT-positive sites, (potential inhibitory synapses), in *Jnk1-/-* neurons (Fig. 6D-F). These results suggest that JNK1 suppresses surface GABAAR-β3 expression, and moreover that it may divert β3-containing GABAARs away from glutamatergic synapses and away from the extrasynaptic sites to inhibitory GABAergic synapses.

**Figure. 6.**
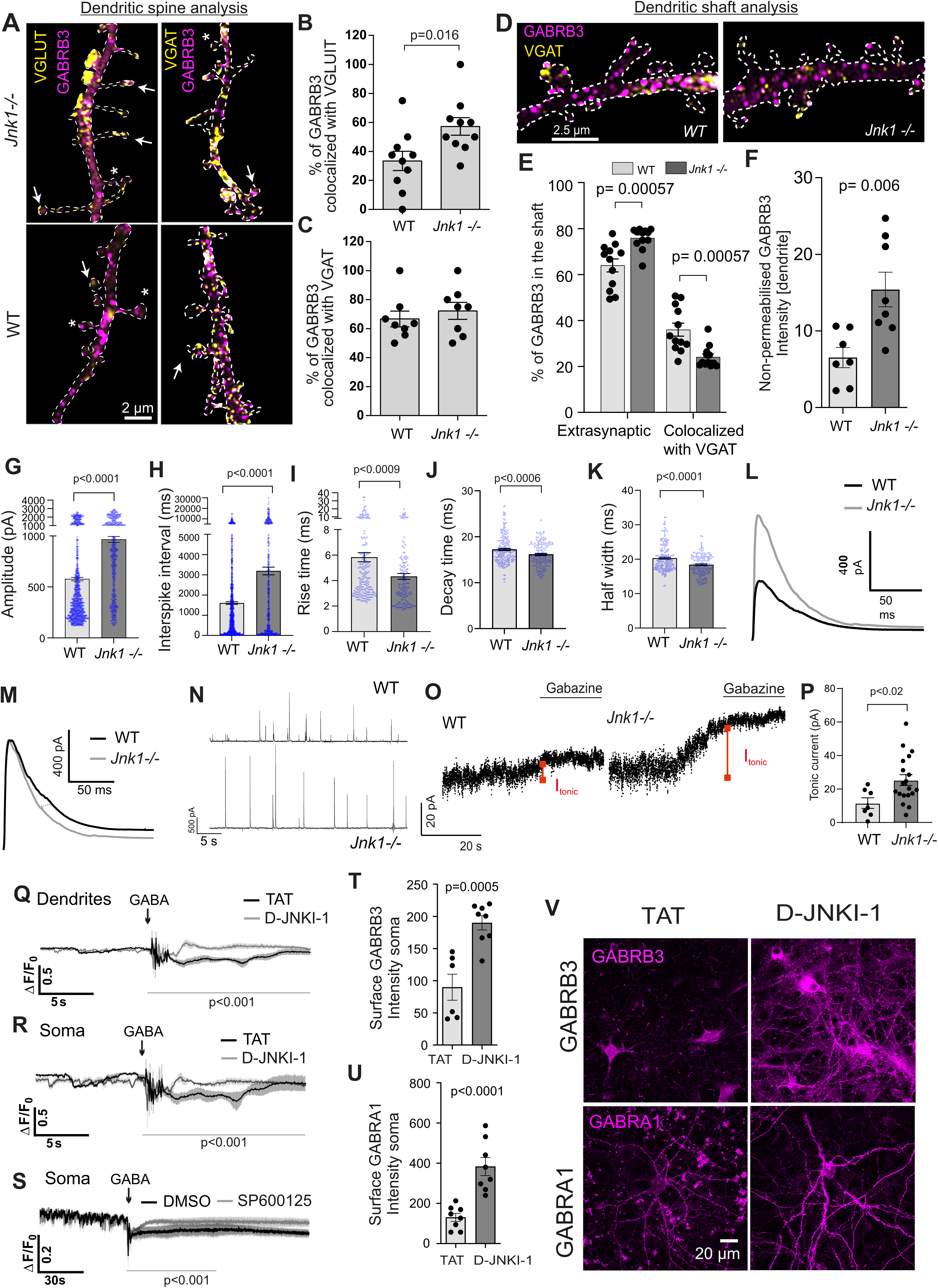
JNK1 decreases spontaneous IPSC amplitude and inhibitory tonic currents and depolarises the membrane potential in dendrites and at the soma. **A.** Representative confocal image of GABRB3 and VGLUT1 or VGAT colocalization in 16 DIV WT and *Jnk1-/-* hippocampal neurons in non-permeabilized cells. The cell outline depicted with a dotted line was traced from co-expressed CFP which acted as a filler. Arrows point to examples where GABRB3 colocalized with VGAT or VGLUT1 while * indicates GABRB3 that does not colocalise with VGAT or VGLUT1. **B-C.** Mean data +/- S.E.M.s from 193 spines in WT and 162 spines in *Jnk1-/-* neurons taken from 3D micrographs from multiple fields in 10 WT and 8 knockout hippocampal neurons. P-value was calculated from Student’s t-test. **D.** Representative images of GABRB3 and VGAT surface staining from 16 DIV WT and *Jnk1-/-* hippocampal neurons. **E.** Quantitative data shows the % of GABRB3 punctae that were extrasynaptic or synaptic. Approximately 200 punctae were analysed per condition. 3D images were taken from 12 r.o.i.s from 8 neurons per condition. P-values were calculated from Student’s t-test. **F** Quantitative data shows total surface GABRB3 levels in the dendritic shaft. **G, H.** Spontaneous inhibitory postsynaptic current (sIPSC) amplitudes were higher in *Jnk1-/-* than in WT neurons, as was interspike interval. **I-K.** Average sIPSCs from *Jnk1-/-* neurons showed faster rise and decay kinetics. Data are shown as mean bars ± S.E.M. for n=3 in both groups. For G-K, statistical significance was between genotypes was tested using unpaired *t*-test. **L.** Average sIPSC traces for single neurons from WT and *Jnk1-/-* groups demonstrate the larger amplitude in *Jnk1-/-* neurons. **M.** After normalization by amplitude, average sIPSC traces from single neurons from WT and *Jnk1-/-* groups demonstrate faster kinetics in *Jnk1-/-* neurons. **N.** Representative traces of outward sIPSCs recorded at -40mV with low intracellular chloride from WT and *Jnk1-/-* cultures are shown demonstrating larger interspike interval and amplitude in *Jnk1-/-* neurons. **O.** Representative traces from tonic current measurements show a shift in inward holding current for the cultured WT and *Jnk1-/-* hippocampal neurons in response to inhibition of extrasynaptic GABAARs with gabazine (250 µM). The time of gabazine action is indicated. Red dots depict the amplitude of the holding current at steady state before and after gabazine. Red lines depict the amplitude of the tonic current (I_tonic_). **P.** Individual tonic current amplitudes are shown. Mean data ± S.E.M. for n=7 for the WT group and n=19 for the *Jnk1-/-* group. Statistical analysis was done with unpaired *t*-test. **Q-R.** Voltron reporter output (meaned) shows average relative transmembrane voltage measurements +/- S.E.M. from control (TAT) and JNK inhibited (D-JNKI-1) hippocampal neurons at 15 DIV following transduction with pAAV-CAG-Voltron/JF549. TAT and D-JNKI-1 (20 µM) were added at -21 h followed by GABA (200 µM) where indicated. Q shows measurements from dendrites (n=8) and R shows measurements from the soma (n=4). P-values were calculated with Wilcoxon test for response period 15s post-treatment. **S.** Comparison between the average control (DMSO) and JNK inhibited (SP600125) intensity traces of Voltron reporter (pAAV-CAG-Voltron/JF549) in hippocampal neurons DIV17, as well as individual traces. DMSO and SP600125 (10 µM) treatment for 21 h, GABA 200 µM. Soma (n=4). P-values were calculated using Wilcoxon test during 60s post-treatment. **T-U.** Quantitative data shows levels of GABRB3 and GABRA1 at the soma of non-permeabilized hippocampal neurons at DIV17 analyzed from confocal micrographs (V). Mean data +/- S.E.M.s is shown. P-values were calculated from Student’s t-test. **V**). Representative maximum projection images of GABAAR-β3 or GABAAR-α1 from the data shown in T-U.

### sIPSCs and tonic GABA current are increased in hippocampal neurons from *Jnk1-/-* mice

We next measured electrical currents in these neurons. Increased expression of cell surface β3-containing GABAARs ought to increase the inhibitory transmission (Jacob et al., 2008). We therefore recorded spontaneous inhibitory postsynaptic currents (sIPSCs). Spontaneous IPSCs solely driven by the baseline activity within the neuronal culture were recorded using low-chloride intracellular solution and voltage clamped at -40 mV. Under these conditions, GABAergic currents appear as outward deflections, while excitatory currents remain inward, allowing straightforward separation by polarity. Cultured hippocampal neurons from *Jnk1^-/-^* mice showed larger sIPSC amplitudes (965±29 pA vs. 578±17 pA, (n=3), p<0.0001) and larger interspike intervals (3191±191 ms vs. 1602±62 ms) (Fig. 6G-I), compared to WT neurons (Fig. 6G-H), suggesting hyperfunction of GABAARs containing β3 (and α1) at the postsynaptic site. Kinetically, iIPSCs from *Jnk1-/-^-^*neurons had sharper rise (4.3±0.2 ms vs. 5.8±0.4 ms, p=0.0009) and decay (16.2±0.2 ms vs. 17.2±0.2 ms, p=0.0006) times, and narrower half-widths (18.4±0.2 ms vs. 20.3±0.3 ms, (n=3), p<0.0001), than those from WT neurons (Fig. 6I-M). Example traces of spontaneous IPSCs demonstrate that the amplitude was higher and frequency decreased in *Jnk1-/-* cultures compared to WT (Fig. 6N). The decreased sIPSC frequency is consistent with decreased number of VGAT-colocalized β3-containing GABAARs and the increased amplitude suggests altered channel properties or subunit composition.

To assess the activity of extrasynaptic GABAARs in hippocampal cultured neurons, we measured inward holding currents before and after application of the GABAAR antagonist gabazine (250 µM). Neurons derived from *Jnk1^-/-^* mice displayed larger tonic currents compared to wild-type neurons (25.4 ± 3.2 pA (n=19) vs. 11.6 ± 3.2 pA (n=7), p=0.02) (Fig. 6 O, P), consistent with increased number or hyperfunction of extrasynaptic GABAARs. Together these findings indicate that in *Jnk1-/-* neurons, both tonic and phasic GABA currents increase, suggesting that JNK1 activity suppresses inhibitory currents.

To cross validate these findings using an orthogonal approach, we used the genetically engineered, membrane-spanning voltage-sensitive reporter Voltron that undergoes FRET when coupled with a fluorescent halotag ligand (Abdelfattah et al., 2019). Membrane potential hyperpolarization is detected as increased signal due to unquenching of fluorescence. Voltron-expressing neurons were treated with TAT or DJNKI-1 for 8 hours and fluorescence was monitored before and after treatment with GABA. While WT neurons underwent a mild hyperpolarisation following GABA, JNK inhibitor-treated neurons showed substantially increased hyperpolarisation, consistent with increased surface GABAAR expression. This response was observed both in dendrites and at the soma with the two independent JNK inhibitors (Fig. 6Q-S). Consistent with this, JNK1 inhibitor treatment increased surface expression of β3- and α1-containing GABAARs at the soma (Fig. 6 T-V), as had already observed in dendrites (Fig. 2).

Together our data uncovers a regulatory role for JNK1 in regulating β3-GABAAR surface expression via a mechanism involving direct phosphorylation of the GIT1 scaffold (summarized in Fig. 7). We propose that JNK1 phosphorylation of GIT1 on S371 disrupts its binding to 14-3-3ζ and paxillin resulting in removal of the β3-containing GABAAR from its complex at the cell surface. Consequently, GABAAR clustering at the cell surface is reduced and GIT1 localisation to dendrites increased where it is bound to β-PIX. This is the first description of JNK1 regulation of GABAAR trafficking and function.

**Figure 7.**
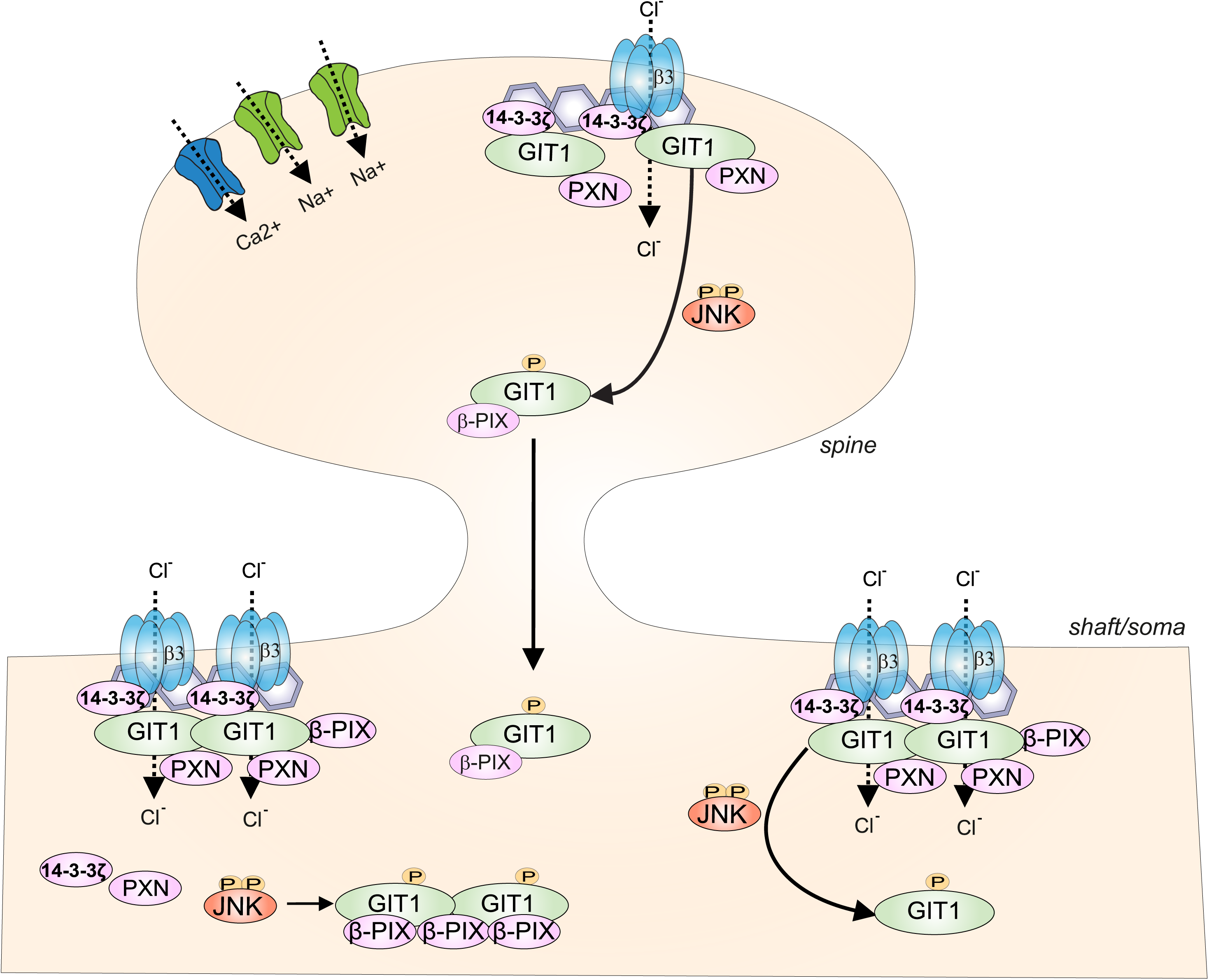
Model depicting JNK1 control of GABAAR surface expression. Active JNK1 phosphorylates GIT1 at serine 371, resulting in reduced surface expression of GABAARs pools localizing at the plasma membrane in dendritic spines i.e. glutamatergic synapses and at extrasynaptic plasma membrane sites in the dendrite and somas. In contrast, inhibition of JNK signaling or genetic deletion of *Jnk1* promotes the association of GIT1 with a plasma membrane-localized receptor complex containing paxillin and 14-3-3ζ proteins. Phosphorylation of GIT1 at S371 disrupts this interaction, leading to dissociation of GIT1 from the complex and subsequence run down of GABAAR surface levels. At the functional level, JNK1 controls spontaneous inhibitory postsynaptic currents (sIPSCs) and tonic inhibitory current. In this way JNK1 may promote neuronal excitability by repressing surface expression of β3-containing GABAARs via a mechanism involving GIT1.

## DISCUSSION

GIT1 has emerged as a scaffolding protein that stabilizes several receptors at the cell surface, including the beta-adrenergic receptor, AMPAR and GABAAR (Premont et al., 1998; Kim et al., 2003; Smith et al., 2014). Genetic variants link GIT1 to attention deficit disorder and schizophrenia (Won et al., 2011; Kim et al., 2017a; Kim et al., 2017b), but how its receptor-clustering function is regulated has been unclear. We show that in the brain, JNK phosphorylates GIT1 on S371, preventing GIT1 from stabilizing β3-containing GABAARs at the cell surface. Accordingly, inhibiting or deleting JNK1 causes GIT1 to accumulate in dendritic spines and increases surface β3-containing GABAARs tethered to paxillin and 14-3-3ζ. This enhances tonic and phasic GABA currents and hyperpolarizes dendritic and somatic membrane potential. JNK also suppresses α1-containing GABAAR surface expression. Thus, JNK controls surface availability of β3- and α1-containing GABAARs, providing a previously unrecognized mechanism for regulating inhibitory signalling.

GABAAR expression at glutamatergic spines is increasingly recognized as a mechanism for fine-tuning excitatory synaptic integration, spine Ca²⁺ dynamics, and plasticity, independent of classical GABA co-release (Hayama et al., 2013; Chapman et al., 2022). We identify that JNK regulates α1- and β3-containing GABAAR localisation at the cell surface in dendritic shafts, in dendritic spines and at the cell soma. GABAARs in spines in turn have been shown to be activated by spillover from nearby inhibitory terminals (Kwon, 2018), representing a shunting mechanism whereby chloride influx modulates glutamatergic synapse excitability. While we do not directly assess GABA/glutamate co-release or whether dual synapses co-exist within a single spine, we do observe that JNK activity depletes the pool of GABAARs in spines that colocalise with VGLUT1, but not with VGAT. Our data shows that JNK activity removes surface GABAARs from spines and dendrites via a mechanism invovlving GIT1, meaning that JNK1 limits inhibitory tone. Analyzing GABAARs at glutamatergic spines is directly relevant to understanding how GIT1-dependent trafficking mechanisms shape the inhibitory/excitatory balance at the level of individual synapses. Although GABAARs at excitatory synapses are often overlooked, several ultrastructural studies have detected β2 and β3 subunits at asymmetric postsynaptic densities alongside glutamate receptors (Knott et al., 2002; Zander et al., 2010; Marlin and Carter, 2014; Vaaga et al., 2014; Kwon et al., 2019). Moreover, GIT1, a known GABAAR scaffold, has been shown to localise in dendritic spines (Zhang et al., 2003). Thus, combined with evidence for glutamate/GABA co-release (Nusser et al., 1998; Fattorini et al., 2009; Marlin and Carter, 2014; Vaaga et al., 2014), our findings support the presence of inhibitory receptors within spines and suggest a role for physiologically active JNK in controlling this localisation.

β3-containing GABAARs negatively regulate spine maturation (Jacob et al., 2009), providing a potential mechanistic link to JNK1-dependent control of spine maturation. In *Jnk1-/-* mice, reduced mushroom spines in CA3 are accompanied by impaired learning (Komulainen et al., 2020), consistent with increased spine-localised β3-mediated shunting inhibition limiting synapse excitability and subsequent spine stabilization. Together, these findings suggest that JNK1-dependent trafficking of β3-containing GABAARs may serve a key role in synaptic plasticity. Consistent with this, GABAAR shunt inhibition in spines is proposed to promote the competitive selection and pruning of spines, as opposed to simply reducing overall neuronal firing (Hayama et al., 2013).

In the present study, the detailed αβγ/δ subunit composition of GABAAR that incorporates β3 and α1 subunits is not known. Multiple GABAAR assemblies containing these subunits have been described across the brain (Olsen and Sieghart, 2009; Ghit et al., 2021). Specifically, β3 and α1 subunits contribute to fast phasic inhibition, where α1β1-3γ2 complexes predominate (Fritschy and Mohler, 1995; Ghit et al., 2021). However, β3 subunit can also regulate inhibitory tone at extrasynaptic sites (Rosas-Arellano, 2025). Some tonic inhibition arises also from β3 homomers and αβ receptors (Mortensen and Smart, 2006), which are constitutively active and poorly inhibited by gabazine (Wlodarczyk et al., 2013); and thus not examined here.

Stabilization of the GABAARs at the membrane does not itself explain their accumulation there; a net increase requires enhanced receptor delivery, such as lateral diffusion, or reduced endocytosis. To test whether endocytosis contributes, we compared the effects of JNK inhibition in cells expressing dynamin-2-K44, a dominant negative dynamin that blocks endocytosis (Ochoa et al., 2000). JNK inhibitor treatment increased receptor surface expression to the same extent as dynamin-2-K44, and combining the two produced no additional effect. This suggests that the mechanism described here may suppress endocytosis, to a similar extent as dominant negative dynamin. Supporting this idea, GIT1 interacts with the endocytosis machinery adaptor AP-2 through residues 300 to 400 and promotes clathrin-mediated endocytosis (Meng et al., 2002; Bai et al., 2011). Thus JNK-dependent regulation of GIT1 may influence receptor surface levels in part by modulating endocytic trafficking.

14-3-3 proteins are conserved scaffolds that bind diverse signaling partners and coordinate their functions (Yaffe, 2002). Several studies show that JNK directly phosphorylates 14-3-3ζ leading to the release of its bound clients (Tsuruta et al., 2004; Sunayama et al., 2005; Yoshida et al., 2005). Here we demonstrate that 14-3-3ζ client binding is required for DJNKI-1-induced upregulation of GABAAR at the cell surface, as the R18 inhibitor of client binding, abolishes this effect. We further show that GIT1-S371A co-immunoprecipitates with 14-3-3ζ, whereas the phosphomimetic GIT1-S371D does not, indicating that the 14-3-3ζ association is necessary for the increase in GABAARs within spines. Conversely, GIT1-S371D preferentially associates with β-PIX, which like GIT1-S371D, translocates out of spines upon JNK inhibition. Together these findings support a model in which GIT1 and 14-3-3ζ act as key components of a JNK-regulated pathway controlling GABAAR availability at the plasma membrane.

We show that neurons from *Jnk1-/-* mice, as well as neurons treated with JNK inhibitors, display markedly increased surface expression of a1- and b3-containing GABAARs. While the physiologically active JNK1 studied here results primarily from synaptic activity (Brecht et al., 2005; Vieira et al., 2010), all JNKs (1, 2 and 3) are activated by stress (Coffey et al., 2002; Coffey, 2014). Our findings indicate that JNK activation reduces GABAAR levels in the plasma membrane resulting in lower inhibitory tone. This ought to prolong stress signaling via the hypothalamus-pituitary-adrenal (HPA) axis by weakening GABA-mediated negative feedback. Consistent with this, deletion of GABAAR subunits induces HPA axis hyperactivity, and anxiogenic and depressive-like behaviors (Shen et al., 2010; Levy and Tasker, 2012; Arora et al., 2024). Similarly, a1-subunit knockout mice exhibit heightened anxiety (Zhang et al., 2010). In post-traumatic stress and panic disorders, GABAAR surface levels are decreased (Bremner et al., 2000; Geuze et al., 2008), and impaired GABAergic signaling contributes to excitatory/inhibitory imbalance and abnormal oscillations in schizophrenia (Uhlhaas and Singer, 2010; Fogaça and Duman, 2019).

JNK activity itself promotes anxiety- and depressive-like behaviors in rodents (Mohammad et al., 2018; Stefanoska et al., 2018), and is activated by stressors relevant to these pathologies, including endocrine and excitotoxic stress (Mukherjee et al., 1999; Qi et al., 2005; Adzic et al., 2009; Kv et al., 2018). Human genetic studies further link JNK-pathway genes to schizophrenia and bipolar disorder (Winchester et al., 2012; Steinberg et al., 2014; Winchester et al., 2014; Marchisella et al., 2016). Thus, identifying JNK as a direct regulator of GABAAR surface expression has broad implications for understanding stress-related neuropsychiatric disorders.

Collectively, our data show that JNK phosphorylates GIT1 to destabilize GABAAR expression at the plasma membrane. This provides neurons with a mechanism to downregulate GABAergic inhibition in response to JNK-activating cues, a process that may contribute to JNK-mediated anxiety behaviours.

## LEGENDS

**Supplementary figure 1. A.** Representative fluorescence micrographs of 16 DIV neurons expressing GIT1-WT, or GIT1-S371A or GIT1-S371D variants are shown (green). The relative expression can be seen in dendrites and spines in WT and *Jnk1-/-* neurons co-expressing mRuby-Lifeact (magenta). Representative maximum projection images are from 25 x optical slices, z = 0.173 µm. **B.** The effect of *Jnk1* deletion and DJNKI-1-treatment on GABAAR surface expression in 16 DIV neurons. Maximum projection images are shown for +\- 20 µM TAT or DJNKI-1 treatment for 8 h. Rescue experiments were carried out by co-transfecting EGFP-GIT1-S371A or - S371D mutants into neurons expressing NT or GIT1 shRNA. **C.** The effect of DJNKI-1 on surface GABAAR enrichment in spines relative to dendrites in non-permeabilized neurons is shown. **D.** Enrichment of total GABAAR in spines relative to dendrites in permeabilized neurons. **E, F.** Raw intensities of surface GABAAR expression in spines and dendritic shafts are shown. Cell surface GABAAR intensities were quantified from multiple images. DJNKI-1 treatment for 8 h increased GABAAR at the cell surface in a GIT1-dependent manner. Mean data +/- S.E.M from ∼40 spines per condition. **G.** Enrichment of total GABAAR in spines relative to dendrites in WT and *Jnk1-/-*transfected with indicated shRNA and GFP-GIT1 variants. **H** Representative images of GABA(A) a1 staining from non-permeabilized 16 DIV hippocampal neurons. Maximum projection images are shown for +\- 20 µM TAT or DJNKI-1 treatment for 8 h. **I, J** Quantitative data showing density of GABA(A)a1 puncta at spine or shaft for neurons in H. Mean data +/- S.E.M from 3-5 cells per condition. **K** Representative images of GABRB3 and VGAT staining from non-permeabilized 16 DIV hippocampal neurons.

## Acknowledgements

This project was funded by grants from the Marie Skłodowska-Curie Actions r’BIRTH Initial Training Network grant #608346 and the Research Council of Finland grant #310583 and #362838 to E.C. We thank the Cell Imaging and Cytometry core which is part of the BioCenter Finland, Finnish Advanced Microscopy Node of Euro-BioImaging Finland, funded by the Research Council of Finland (FIRI grants #359073, #358879) for providing infrastructure support.

