## Supplementary figures and images for "JNK regulates β3-containing GABAAR expression at the cell surface via the receptor clustering protein GIT1 (ArfGAP1)"

### Supplementary figure 1

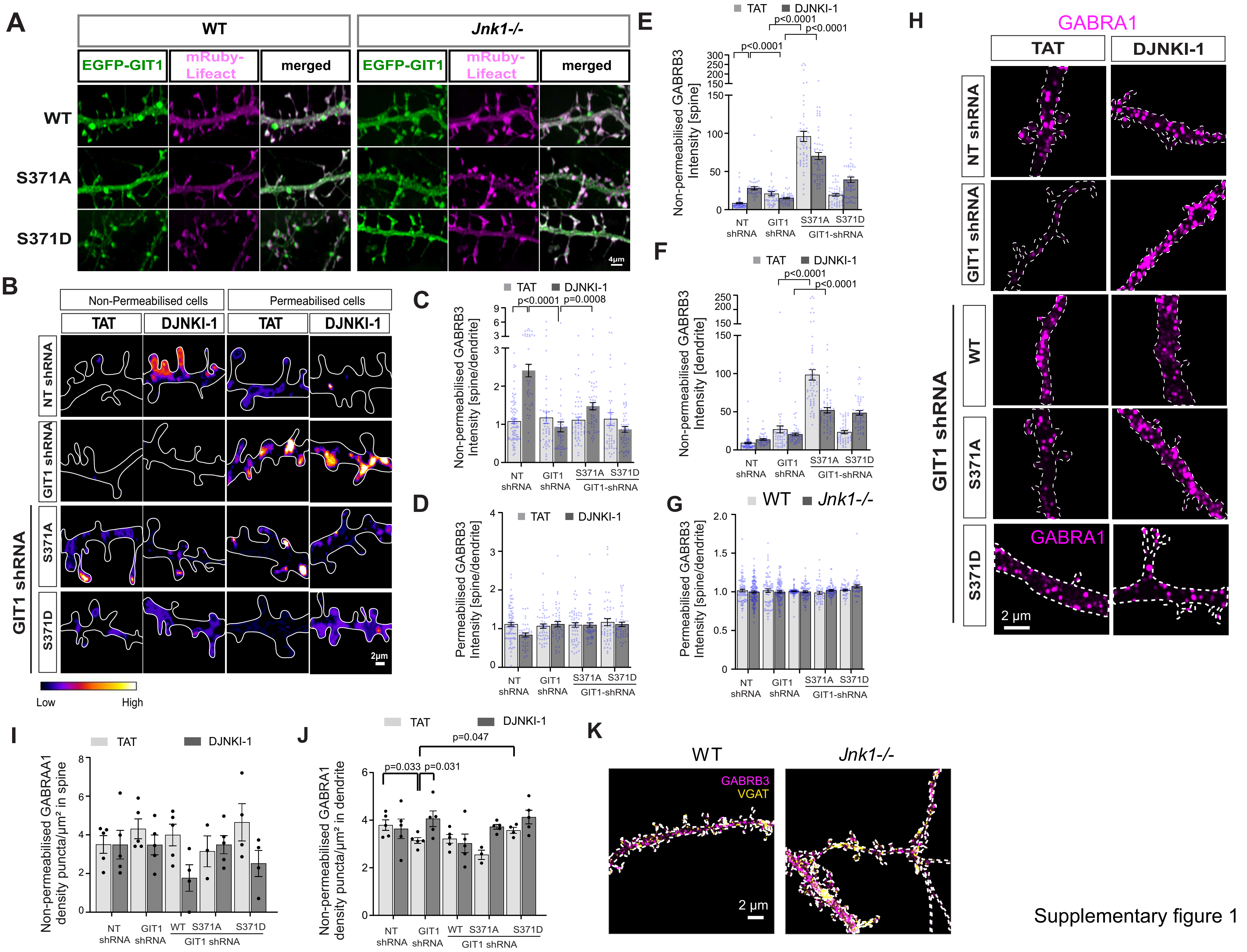


Supplementary figure 1
